# The neural correlates of parallel and serial search

**DOI:** 10.64898/2025.12.29.696840

**Authors:** G. Kandemir, D. H. Duncan, D. van Moorselaar, J. Theeuwes

## Abstract

For almost half a century, target-distractor similarity has been known to induce different visual search modes. When a target is highly salient, it can pop out, suggesting parallel processing of all items irrespective of set size. By contrast, high similarity among items requires item-by-item comparison with an attentional template, a characteristic of serial search. Despite this long-standing distinction, little is known about the neural correlates of search modes, as typical differences in visual displays confound interpretation. Here, we contrasted the neural correlates of serial and parallel search under visually identical displays. Across distinct blocks, we biased 24 participants (21 female) toward parallel or serial search by varying target-distractor similarity, thereby directing attentional focus toward a single feature or conjunction. Embedded among inducer trials, test trials were visually identical across all blocks and afforded both parallel and serial search. Behavioral analyses confirmed successful induction of distinct search modes during test trials. EEG decoding reliably discriminated search modes and these neural patterns generalized across inducer and test trials. Attentional deployment toward the target differed across search modes, revealing topographical differences in target location representations. The strength of target-location representations correlated with response times, indicating that during parallel test trials participants switched search strategies when the target was not detected early. Moreover, target representations diverged between search modes: a temporally stable pattern emerged during serial search, suggesting reliance on working memory, whereas parallel search was characterized by more dynamic representations, likely reflecting prioritization of the relevant feature. These findings demonstrate that search history shapes search mode, giving rise to clearly distinct neural dynamics even under visually identical stimulation.

## Introduction

Searching for an apple differs qualitatively depending on whether it appears among oranges or among peaches: in the former case it can be detected “preattentively,” via parallel processing across the display, whereas in the latter it requires focal attention to be deployed serially from item to item or small groups of items. (Neisser, 1967). Feature Integration Theory (FIT) (Treisman & Gelade, 1980) lies at the core of the debate about when visual search proceeds in parallel and when it is carried out serially. FIT originally distinguished between preattentive and attentive search. Preattentive processing was proposed to operate in parallel across the visual field and to be limited to basic features such as color, motion, size, and orientation. Targets defined by a single feature (e.g., a red item among green) could be detected by parallel search yielding reaction times largely independent of display size (Bergen & Julesz, 1983; Treisman & Gelade, 1980; Theeuwes, 1992). In contrast, attentive processing was required to bind features together to identify complex objects (e.g., a T among Ls). This process was assumed to operate serially, item by item, producing reaction times that increased with set size, typically by 20–40 ms per item (Wolfe et al., 1989; Wolfe, 2020).

For the last half century, the distinction between parallel and serial search has shaped much of the psychophysical literature and lies at the core of classic attentional theories, including Feature Integration Theory (Treisman & Gelade, 1980), Guided Search (Wolfe et al.,1989), Feature Similarity (Duncan & Humphreys, 1989), Biased Competition Theory (Desimone & Duncan, 1995) and TVA (Bundesen, 1990). This work has relied heavily on the behavioral observation of search slopes – where the mean reaction time increases reliably in serial search as the number of items increase, but not in parallel search paradigms (Treisman & Gelade, 1980).

Although the parallel–serial distinction features prominently in most theories of visual search, treating search slopes as a straightforward diagnostic is problematic. For instance, a “serial” slope of 20–40 ms per item would imply that target–template matching can be completed within a few tens of milliseconds which is far faster than any plausible object-recognition mechanism (Thorpe et al., 1996). The problem becomes even clearer when other measures of attentional dwell times are used. For example, during visual search Theeuwes et al (2004) showed dwell times during visual search of about 250 msec/item, a number comparable to dwell time found in paradigms such as the attentional blink (Raymond et al., 1992; Duncan et al., 1994).

One response to this problem has been to argue that visual search may not necessarily have a serial component. Instead, object recognition may rely on parallel processes capable of evaluating many items simultaneously. Several forms of parallel processing can reproduce the reaction-time patterns that originally motivated serial, self-terminating search models (Palmer, 1994). These results have led to the view that visual search may not involve two qualitatively distinct modes. Rather, differences in search efficiency may reflect how display density affects the discriminability of targets and distractors (Eckstein, 1998; Pashler, 1987; Palmer, 1994; 1995).

Related to these issues is the notion that increasing the number of items in a display reduces inter-item distance, thereby increasing local contrast interactions and lateral masking (Wang & Theeuwes, 2020). These perceptual factors can alter target–distractor discriminability and produce reaction-time differences independent of the underlying search mode. Moreover, as argued by Theeuwes (2004), flat set-size functions do not necessarily imply truly parallel search. Instead, search may proceed serially over small groups of items, with limited parallel processing within each group—a clump-wise search process (de Waard & Theeuwes, 2026; Liesefeld & Müller, 2020; Theeuwes, 2023a, 2023b).

Because the parallel–serial distinction is central to theories of visual search and interpretations based on search slopes remain controversial, it is essential to determine whether these putative search modes differ at the neural level. The present study aimed to demonstrate distinct search modes by identifying clear neural dissociations between serial and parallel search, while carefully controlling for differences in the search displays themselves. Such a neural comparison allows the debate to move beyond behavioral measures based on search slopes and would reveal the neural mechanisms underlying these putative search modes. To this end, we built on recent evidence showing that selection history strongly influences attentional selection (see Theeuwes et al., 2023; Theeuwes, 2025). Using so-called inducer trials, participants were biased toward a search mode promoting either serial or parallel processing. Subsequent critical test trials were then used to assess the neural signatures associated with each induced search mode..

In the current study, participants had to search for a rotated target T among Ls and report whether the T was titled to the right or left. To induce serial search, we manipulated shape and orientation similarity, by extending short arm of distractor ‘L’s to make them appear similar to target ‘T’ and orienting all distractors heterogeneously in cardinal directions. For parallel search inducers, the nontarget elements were lowercase L’s, orientated unanimous in cardinal directions. During test trials, the target remained the same yet the nontargets consisted of uppercase letter ‘L’ with long arm vertically aligned (see Figure 1). We hypothesized that differences in search modes would lead to distinct neural patterns, distinguishable by multivariate analysis of the EEG. Our focus was on how search type affected the modulation of target location and relevant feature processing.

**Figure 1.**
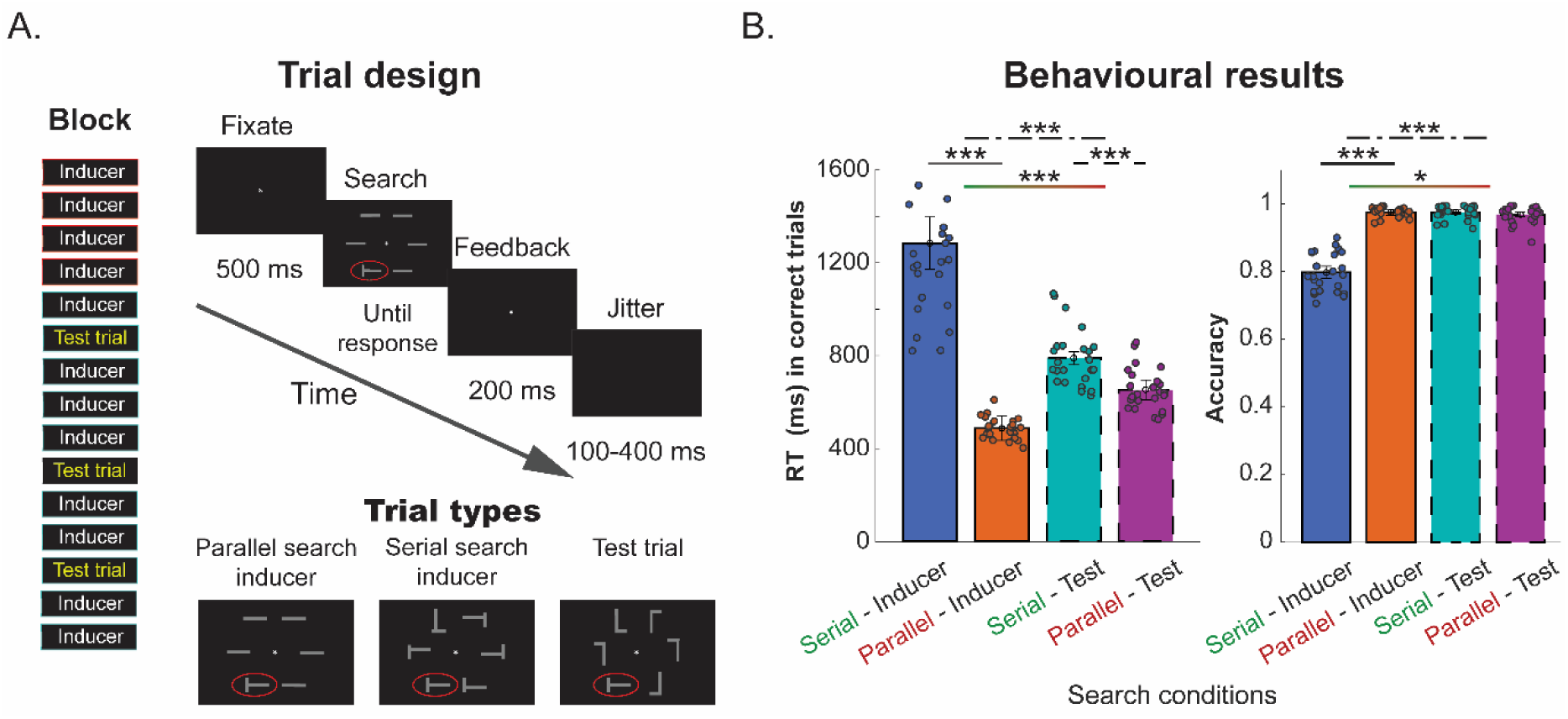
Trial design (A) and behavioural results (B). Note. Trial design and behavioral output. **A.** A trial consisting of the presentation of a fixation dot, followed by search display until subjects reported tilt direction of target (T), followed by feedback and intertrial interval. The similarity of distractors to target was varied with regards to shape and orientation across parallel and serial inducers as well as test trials. The block order reflects that initial four trials (with red borders) were always inducers of one kind whereas three test trials were embedded randomly amongst subsequent 8 inducers (blue borders) of the same kind, where search type was swapped in subsequent blocks. **B.** Bargraphs showing mean RT (left) in correct trials under distinct search conditions and accuracy (right) overall. The colored dots reflect subject means whereas whiskers show 95% CI for the group mean, corrected for subjects. Asterisks indicate statistically significant comparisons (*, *p* < 0.05; **, *p* < 0.01; ***, *p* < 0.001)

## Results

### Behavioural results

Participants (n= 24, 21 female, M_Age_ = 20.79, s.d._Age_ = 1.50) searched for a T (1.5° x 1.125° visual angle) amongst 5 distractors, all displayed evenly distributed on an invisible circle (7° visual angle in diameter) with an origin at the center of the screen. Blocks were 15 trials long, made up of 12 inducer trials and 3 test trials. Each block used one type of inducer (serial or parallel) with the inducer type swapping between blocks (Figure 1A). In the serial inducer trial, distractors were distorted Ls with shorter edge sticking out from opposite end (1.5° x 1.125° visual angle), presented in different cardinal orientations. In parallel inducer trials, distractors were flat lines, all vertically or horizontally oriented (1.5° x 0.15° visual angle). In test trials, distractors were proper Ls with distinct orientations where all long sides were vertically oriented (1.5° x 1.125° visual angle). In each block, the three test trials were positioned randomly anywhere between fifth and 15^th^ in the sequence, ensuring that the participant began each block with at least four trials of the blocks main inducer type (See figure 1A for the full trial and block schematic).

We investigated response time (RT) in correct trials (see Fig. 1B). The distribution was positively skewed, deviating from normality, therefore log transformed RT was assessed with 2×2 RM-ANOVA with trial type (test vs. inducer) and search type (serial vs. parallel) as main factors. Omnibus test showed that RT varied across trial and search type conditions as the main effects of trial type, (*F*(1,23) = 18.575, MSE = 0.01, *p* < 0.001, *η²* = 0.015), and search type (*F*(1,23) = 530.668, MSE = 0.014, *p* < 0.001, *η²* = 0.638) were significant. The effect of trial condition on RT varied based on search mode as the interaction also reached statistical significance (*F*(1,23) = 403.291, MSE = 0.008, *p* < 0.001, *η²* = 0.285). Additional planned comparisons showed that parallel search was faster than serial search both in inducer and test trials; *t*(23)= 30.05, *p* < 0.001, and; *t*(23)= 6.68, *p* < 0.001, respectively. This latter finding is crucial, as it demonstrates that we successfully induced a particular search mode while keeping the search display identical across the two conditions.

Accuracy was overall high (See Fig. 1B). Accuracy was not normally distributed, therefore logit transformed data was tested with RM-ANOVA where trial type (inducer, test) and search type (serial, parallel) were factors. Both trial type, (*F*(1,23) = 24.082, MSE = 5.885, *p* < 0.001, *η²* = 0.256), and search type (*F*(1,23) = 18.462, MSE = 3.452, *p* = 0.03, *η²* = 0.033) as well as their interaction (*F*(1,23) = 22.251, MSE = 3.954, *p* < 0.001, *η²* = 0.159) were statistically significant. Performance was overall higher in test trials, where the difference between search types was small and statistically not significant, *t*(23) = 1.354, *p* = 0.19. In case of inducers, serial search trials produced lower accuracies than parallel search trials (*t*(23) = 5.026, *p* < 0.001). The results reflected that overall parallel search was easy when target and distractors could be distinguished on a single feature.

## Multivariate analyses

### Neural signatures distinguish parallel and serial search modes

We first asked whether distinct search modes could be discriminated from EEG activity recorded during test trials with identical visual displays. Using 8-fold cross-validated decoding from 17 posterior electrodes, search type (parallel versus serial) was successfully decoded shortly after display onset (p < 0.001, cluster corrected; Fig. 2A, black line). Critically, a control analysis suggested that this effect was not driven by gaze position, which showed no significant correlation with EEG-based decoding across time, trials, or subjects (Fig. S1B).

**Figure 2.**
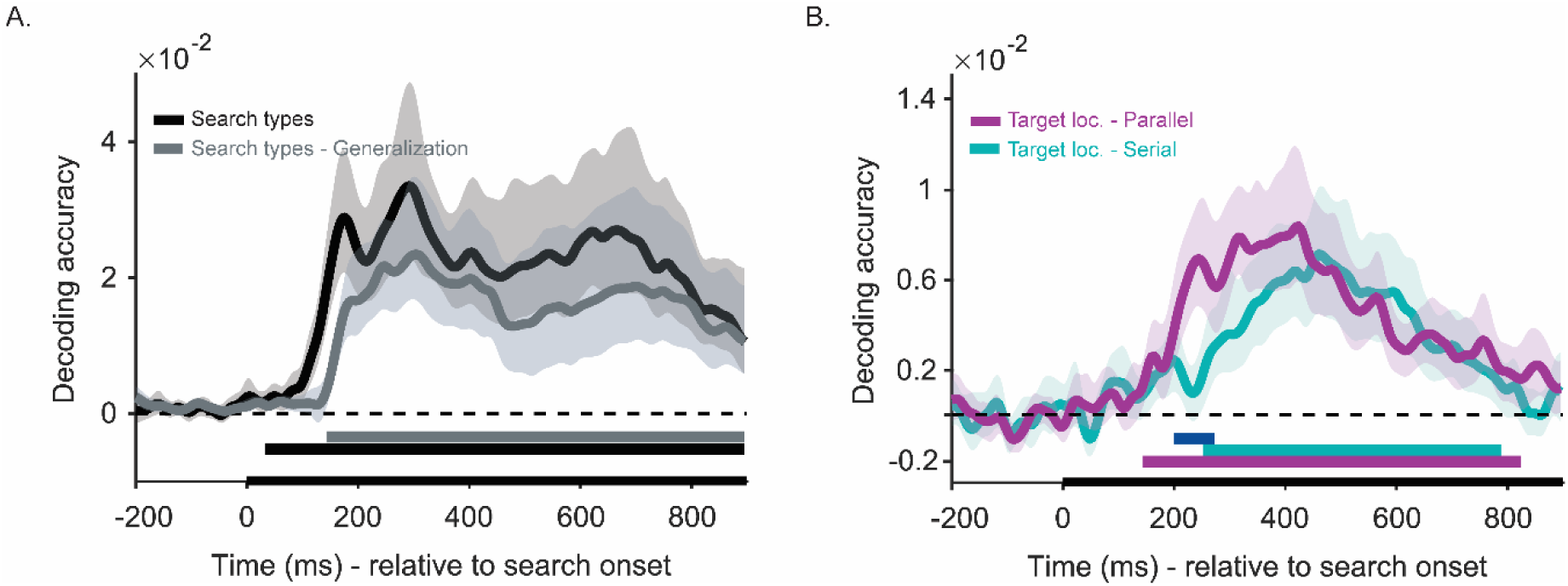
Decoding search types in test trials and the generalization of neural code across trial types as a function of time (A), and decoding of target location in test trials during parallel and serial search (B). *Note.* Decoding search type and target location. **A.** Time-course decoding for Search type (black) and the generalization of neural code when training on inducers and testing on test trials (gray). Solid lines reflect mean decoding accuracy as a function of time. Shades around solid lines mark 95 % C.I. Solid bars below baseline reflect the time-window with statistically significant decoding according to cluster-based permutation test (*p* < 0.05, two-tailed). **B.** Topographic distribution of search type according to searchlight analysis, within three time-windows ranging from 0 to 300, 300 to 600 and 600 to 900, respectively. Brightness reflects the strength of decoding whereas dots (black) mark electrodes with a statistical significance (*p* < 0.05, one-tailed).

To test whether these neural signatures reflected stable differences across search modes, that generalize across distinct visual contexts, we trained decoders on inducer trials and tested on test trials. Despite substantial visual differences between inducers and test displays, decoders successfully generalized (*p* < 0.001, *cluster corrected;* Fig. 2A, grey line), emerging approximately 100 ms later than within-test decoding.

### Target location decoding reveals differential attention deployment

We next decoded target location in test trials to characterize how distinct search modes guide spatial attention on identical search displays. Target location was successfully decoded under both parallel (*p* < 0.001, *cluster corrected; Fig. 2B, violet*) and serial search (*p* < 0.001, *cluster corrected; Fig. 2B, cyan*). Critically, location information emerged earlier and more strongly during parallel search (*p* = 0.049, cluster corrected), consistent with faster behavioral responses and mirrored by the N2pc component Supplements – 3, Figure S3), an established index of attentional selection. As with search type decoding, gaze position was decodable but showed no correlation with EEG-based location decoding (Fig. S2), strongly suggesting that covert attentional shifts, not overt eye movements, drove these effects.

### Cross-generalization reveals mode-specific attention deployment

Having established that neural signatures distinguished between search modes, we investigated whether parallel and serial search engaged fundamentally different patterns of spatial attention. In doing so we first confirmed that inducer trials indeed elicited distinct attentional profiles by decoding target tilt at each location separately. Parallel inducers showed a consistent time course across all locations with clear, overlapping peaks, indicating simultaneous processing. In contrast, serial inducers showed highly variable timing across locations with no consistent peaks, indicating that even items at the same spatial position were attended and processed at different time points throughout trials (Figure S4). This was also confirmed by temporal generalization of target location in inducers which indicated that a general attentional spotlight was used in both search modes, with highly distinct temporal characteristics (Figure S5).

We then trained location decoders on inducer trials and tested on test trials to assess how induced search modes shaped attention deployment. In parallel search test trials, target location was strongly decoded when training on parallel inducers (p < 0.001; Fig. 3A, purple) but only weakly when training on serial inducers (p = 0.002–0.038; Fig. 3A, orange), with significantly higher accuracy for parallel-trained decoders during an early-to-mid latency period (p ≤ 0.002). Importantly, trials with stronger parallel-trained decoding in an early time window (100-300 ms) showed faster response times (t(23) = −4.00, p < 0.001; Fig. 3A inset), indicating that trials with stronger parallel search signatures responded faster whereas failure to find target early probably initiated serial search.

**Figure 3.**
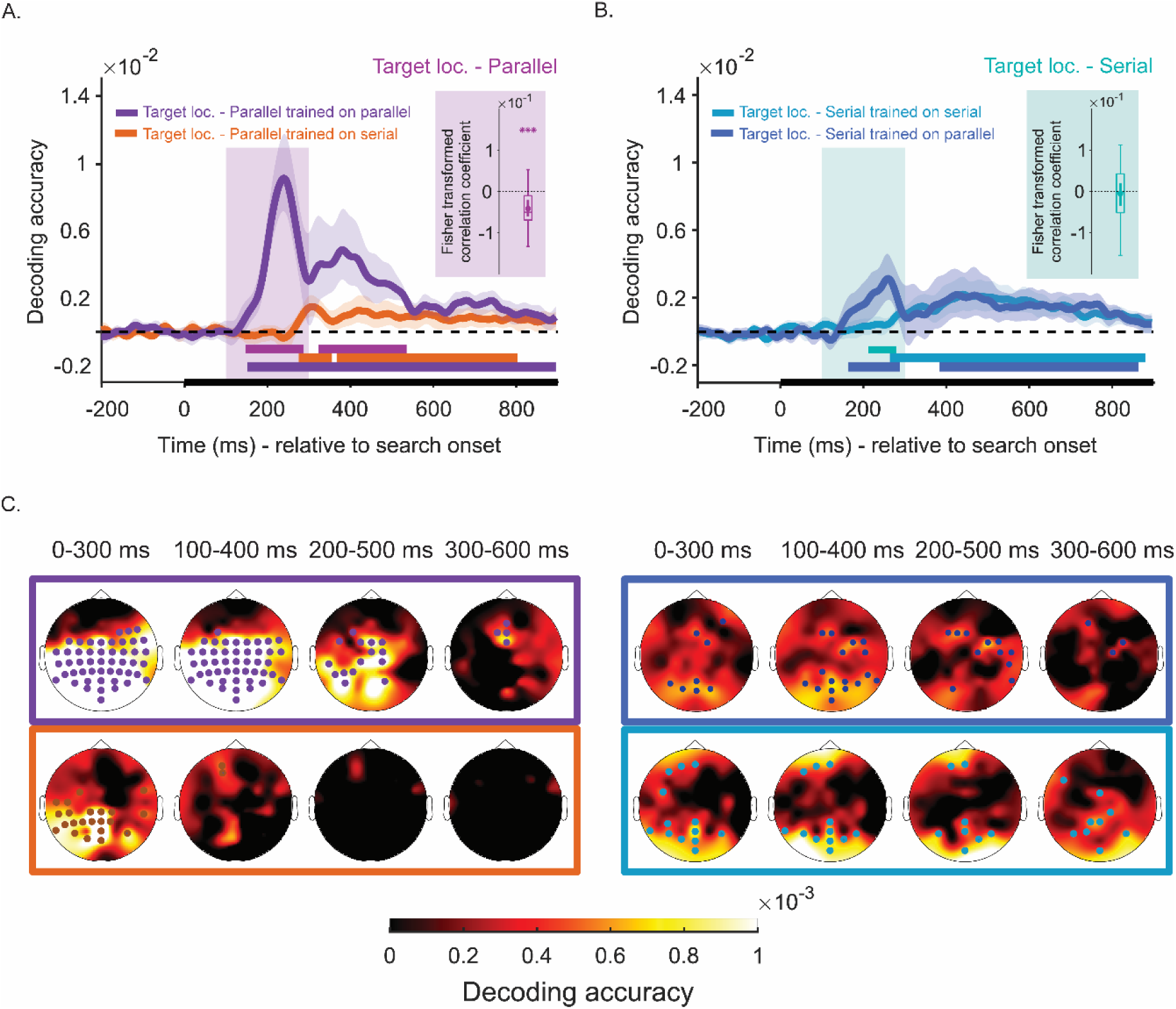
Generalization of target location representation in inducer trials on test trials with parallel (A) and serial search (B),and the topographical distribution of the generalized signal in first 600 ms period in 300 ms wide windows in steps of 100 ms. **(C)** *Note.* Generalization of location modulation between inducer and test trials. **A.** Time-course decoding of target item location in test trials with parallel search when training on inducers with serial search (orange) and parallel search (purple). Boxplot shows Fisher transformed correlation coefficients showing relationship between RT and the decoding accuracy difference within pink colored zone. Asterix indicates mean value significantly differs from zero (***, *p* < 0.001) **B.** Time-course decoding of target item location in test trials with serial search when training on inducers with serial search (teal) and parallel search (blue). Boxplot shows Fisher transformed correlation coefficients showing relationship between RT and the decoding accuracy difference within colored zone. **(A-B)** Solid lines reflect mean decoding accuracy as a function of time, indicating generalization of location modulation across inducer and test trials in specific conditions. Shades around solid lines mark 95 % C.I. Solid bars below baseline mark the time-window with statistically significant decoding according to cluster-based permutation test (*p* < 0.05, two-tailed). **C.** Topographic distribution target location signal during parallel (left) and serial (right) search when training on inducers separately for parallel (top) and serial (bottom) search using searchlight analysis over windows ranging from 0 to 300, 100 to 400, 200 to 500 and 300 to 600 ms, respectively. Brightness reflects the strength of decoding whereas colored dots mark electrodes with statistically significant contribution (*p* < 0.05, one-tailed).

Serial search test trials showed a different pattern. Target location generalized from both serial inducers (p < 0.001; Fig. 3B, teal) and parallel inducers (p ≤ 0.017; Fig. 3B, blue), with parallel-trained decoders showing slightly earlier emergence during an early period (p = 0.04). However, unlike parallel search, decoding strength did not correlate with response time (t(23) = −0.38, p = 0.70), suggesting that even when targets were located early, serial search engaged additional processing stages that delayed responses.

To identify which electrode locations best captured these cross-generalization effects, we conducted searchlight analysis (300 ms sliding windows over the first 600 ms), decoding target location from each electrode and its two nearest neighbours. This revealed distinct topographical signatures between search modes. Parallel search test trials showed sustained posterior parietal involvement when trained on parallel inducers, lasting approximately 500 ms (Fig. 3C, left, purple), but minimal generalization from serial inducers after the initial period (Fig. 3C, left, orange). In contrast, serial search test trials engaged both frontal and occipital electrodes regardless of training condition (Fig. 3C, right), with effects persisting longer than in parallel search. This frontal involvement in serial search and posterior parietal emphasis in parallel search aligns with established neural correlates of these search modes (e.g. Bushman & Miller, 2007), crucially demonstrated here for visually identical displays.

### Target representations differ between search modes

The distinct attentional profiles of parallel and serial search suggest they may also differ in how target features are neurally represented. To investigate this, we decoded target tilt direction using temporal generalization analysis, training and testing decoders at all time points to characterize the stability of target representations. We focused on inducer trials as they provided sufficient trial counts for this analysis. To account for variable target detection timing, we time-locked epochs to response execution (−400 to 0 ms relative to response) and included only trials with RTs between 400–1500 ms to increase temporal consistency (∼2 SD from the group mean corresponding to 80-90% of the data per condition).

During parallel search, target tilt decoding was highly time-specific, with significant clusters aligned along the diagonal (*p* < 0.05, cluster corrected*; black outline in left panel Figure 4*), and diagonal decoding significantly exceeding off-diagonal generalization (p < 0.05, cluster corrected, white borders, *one-tailed*), indicating dynamic representational changes throughout the trial.

**Figure 4.**
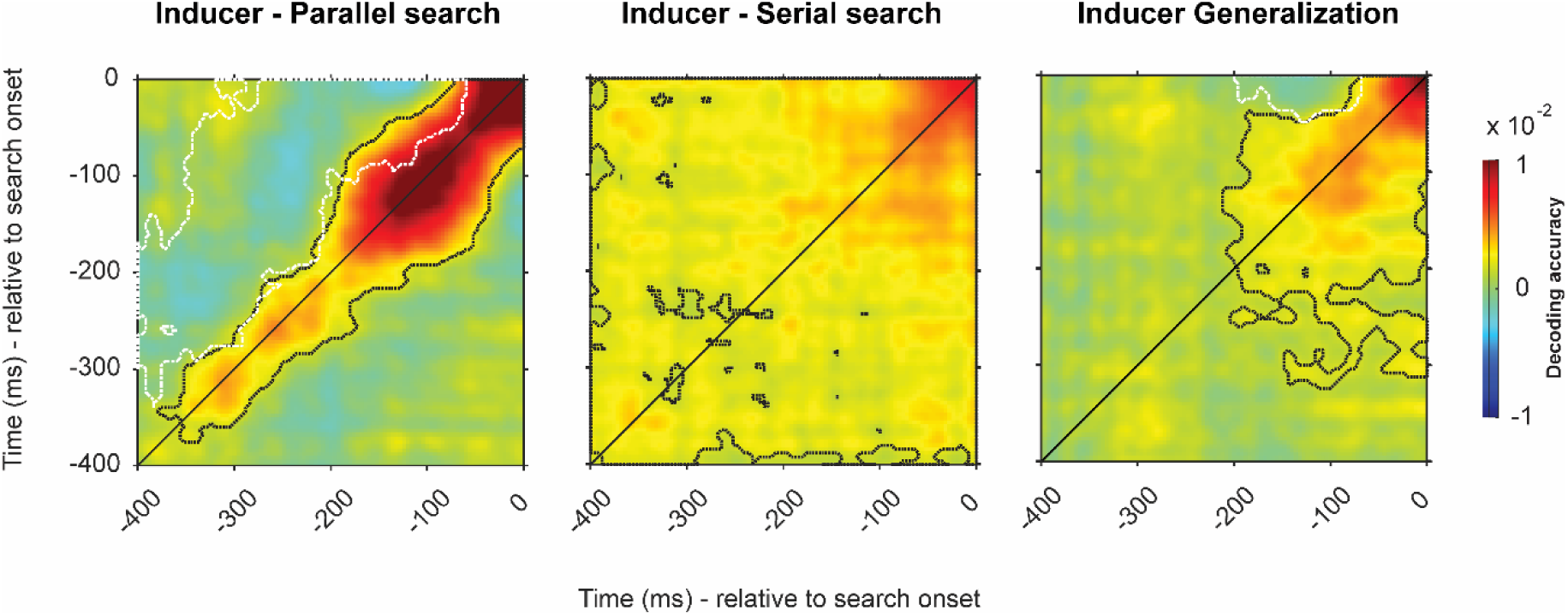
Temporal generalization of neural code for target tilt direction in parallel (left), serial search inducers (middle) and the generalization across (right), relative to response time. *Note.* Decoding target tilt direction by training at a time point and testing on all time points. Temporal generalization is assessed within and across each search type in inducer trials. Dotted black lines mark clusters with statistically significant decoding against zero (*p* < 0.05, two-tailed). Dotted white lines mark clusters with diagonal decoding is significantly higher than the average of vertical and horizontal, indicating dynamic decoding period (*p* < 0.05, one-tailed).

In stark contrast, serial search showed broad temporal generalization across nearly all time points, with no diagonal preference (Fig. 4, middle, *p* < 0.05, *two tailed*), indicating that target representations remained stable throughout the pre-response period. This sustained representation may reflect an attentional template in working memory retrieved from long-term store once the sequential comparison detected the target item, a step that appears unnecessary in parallel search due to rapid extraction of task relevant feature, namely the tilt direction of the target. Cross-generalization from serial to parallel inducers revealed significant overlap only in the final 200 ms before response (*p* < 0.05; Fig. 4, right, black borders), likely reflecting shared response selection and execution mechanisms. The absence of earlier generalization confirms that target feature representations differ fundamentally between search modes.

Together, these results demonstrate that serial and parallel search modes engage qualitatively distinct neural mechanisms: Parallel search relies on rapid, posterior-mediated attention with dynamic, time-varying target representations, whereas serial search involves sustained frontal-posterior networks supporting sequential attention deployment with stable target representations that may reflect template-matching target shape rather than task relevant feature.

## General Discussion

The study investigated neural signatures underlying serial and parallel search, using visually identical test trials embedded amongst inducer trials designed to bias subjects toward a specific search mode in each block. Serial and parallel search was facilitated in separate blocks through manipulations of target–distractor similarity and orientation homogeneity. Embedded within these blocks were test trials that could be effectively completed with either parallel or serial search. Behavioral results showed that participants were strongly biased to use the search mode that was prominent within that block, showing that a specific search mode can be induced by previous trials. This allowed us to then observe the EEG correlates of serial and parallel search in these test trials which were perceptually matched.

Behaviourally, the results indicate that it was indeed possible to induce different search modes using the same visual search displays. In serial search inducer trials, response times were slowest, and accuracy was lowest, clearly differing from parallel search trials. When comparing test trials, we observed a similar pattern in response times: participants responded faster following parallel search induction.

EEG decoding further demonstrated that, induced search mode could be distinguished while searching visually identical displays. This confirmed that qualitatively different cognitive mechanisms were employed depending on whether serial or parallel search was expected on the upcoming trial. The neural patterns distinguishing search modes generalized across inducer trials with distinct visual displays, suggesting that decoding seized upon neural processes recurring in search modes. In line with this, we observed that target location was detected more quickly during parallel search whereas it gradually peaked during serial. The temporal nature of target location overlapped with N2pc component associated with target detection (Eimer, 1996), providing further evidence for differences in the neural signatures of search modes. Considering that induced search modes were frequently altered in consecutive blocks, the effect cannot be explained by temporal autocorrelation.

Serial and parallel search showed clear differences in the deployment of attention during test trials. When decoders were trained on inducer trials and tested on the identical test displays, target location could be decoded in both cases, as by using serial on parallel and vice versa, indicating that spatial information generalized across search modes. However, the timing and the strength of generalization differed, suggesting that serial and parallel search rely on distinct temporal dynamics of attentional deployment even for identical sensory input, while congruency of search modes yielded stronger decoding. Specifically, we observed an early peak in target location generalization when trained with a parallel inducer, but a slower and more delayed onset when training with serial search inducers. These differences in decoding trends could be indicative of search mode mix-ups in a subset of test trials, or alternations if for example parallel search failed. Supporting this, we observed that in parallel search test trials, responses slowed down if generalization from parallel inducer relative to serial inducer dropped, suggesting that if target could not be located early on with parallel search, serial search likely was initiated. On the other hand, in serial search test trials, response speed did not correlate with the difference of target location generalization between search types. This may reflect the fact that during serial search, even when the target is located early on some trials, additional cognitive operations, such as verifying the target through a match with attentional templates, may still be required, thereby delaying responses relative to parallel search. This interpretation would align with search models such as Guided Search (Wolfe, 1994; 2025), in which both top-down and bottom-up signals contribute to attentional selection depending on search mode.

The generalization of target location from inducers to test trials yielded topographical differences across search modes. In parallel search test trials, target location strongly generalized with both types of inducers, mainly reflected in electrodes above posterior parietal regions and occipital cortex. This effect was mainly observable within the early 400 ms time window. In serial search test trials, generalization was evident in occipital but also frontal electrodes, sustained over a longer period relative to parallel search.

The topographical distinctions across search modes align with previous reports. Earlier research associated serial search with the involvement of frontal regions, possibly reflecting spatially guided shifts of attention across items (Anderson et al., 2007; Donner et al., 2002; Leonards et al., 2000; Nobre et al., 2003) and it is assumed that as target salience increases, the demand for top-down control diminishes, allowing attention to be guided more automatically by bottom-up salience maps. Contrasting search types, differential activity was reported in the superior parietal lobule (Corbetta et al., 1995), while the involvement of frontal regions, particularly dorsolateral frontal cortex (DLPFC) and frontal eye fields (FEF), has been more mixed across search types (Corbetta & Shulman, 2002; Iba & Sawaguchi, 2003; Ossandón et al., 2012). FEF activity has been linked to target selection in both search types, potentially driven by both bottom-up salience and top-down control mechanisms (Anderson et al., 2007; Buschman & Miller, 2007; Thompson & Bichot, 2005; Thompson et al., 2005; Purcell et al., 2012). In our study, despite the limited spatial resolution of EEG, we also observed clear distinction in the involvement frontal electrodes during distinct search modes.

Our results revealed an additional difference between search types in how the target feature was represented. During parallel-search inducer trials, the neural code for target tilt was dynamic and changed over time, whereas during serial search it was temporally stable. Notably, these representations did not generalize across search conditions until the final 200 ms before the response, a period likely reflecting response selection and execution. The lack of early generalization, despite within-condition decodability of target tilt, suggests qualitative differences in representational content. In parallel search, a highly salient target is detected rapidly through bottom-up processing, after which top-down selection modulates only the task-relevant feature, namely tilt direction. In contrast, serial search requires prior verification of the target letter among similar distractors, necessitating modulation of the entire shape. This process may involve reactivation of a long-term memory target template (Wolfe, 2012; Cunningham & Wolfe, 2014; Wolfe, 2020) or temporary storage of the full target shape in working memory to protect it from sensory competition until tilt information is extracted. This account is consistent with evidence for temporally stable neural codes in long-term memory representations (Kandemir & Akyürek, 2023) and with findings that working memory preferentially stores task-relevant information—tilt direction in parallel search and full shape in serial search (Bocincova & Johnson, 2019; Serences et al., 2009).

In sum, for more than three decades, the distinction between serial and parallel search has been heavily debated, largely because interpretations based on search slopes are ambiguous and controversial. The present study provides some of the strongest evidence to date for genuine neural differences between serial and parallel search modes that are not confounded by display characteristics. In addition, we show that both processing at the target location and the neural representation of task-relevant target features differ across search conditions, despite identical visual input.

## Methods

### Participants

The sample of 24 subjects (21 female, M_Age_ = 20.79, s.d._Age_ = 1.50) studying at the Vrije Universiteit Amsterdam volunteered to take part in the study in return for monetary reward (10 € per hr) or course credits. An additional 3 participants were removed from the sample due to excessive noise, high eye and muscle artefacts or faulty electrode configurations. The sample size was inferred from earlier studies using the same analyses techniques on visually presented stimuli (e.g., Wolff et al., 2017). The participants were informed on the experimental design, and procedure as well as data sharing and storing practices both verbally and in written format, and a written consent for participation was collected. An ethical approval on the experimental protocol in line with the principles of the declaration of Helsinki was acquired from the Scientific and Ethics Review Board of the Faculty of Behavioral and Movement Sciences at VU University Amsterdam (protocol number VCWE-2021-173)

### Stimuli and Apparatus

All stimuli were created and presented using Psychtoolbox-3 for MATLAB (Brainard, 1997; Kleiner et al., 2007). The experiment was displayed on a 23.8 inch (60,452 cm on diagonal) ASUS ROG Strix XG248Q monitor with a refresh rate of 240 Hz and a resolution of 1920 by 1080 pixels.

Throughout the experiment, a black background was maintained (RGB = 10 10 10). A fixation dot was displayed at all times except for a brief period (Range= 200 – 600 ms duration) between consecutive trials in order to allow blinking. The fixation dot consisted of a small white disc (RGB= 255 255 255; 0.22 degrees of visual angle diameter-dva) superimposed on a black cross (0.33 dva length, 0.13 dva line thickness) and a large white disc (RGB= 255 255 255; 0.66 dva) (Thaler et al. 2013).

The search display contained 6 items (5 distractors and 1 target) displayed in gray (RGB = 128 128 128), and positioned equidistantly on an invisible circle around fixation (7 dva in diameter). The stimuli were located at 30° , 90°, 150°, 210°, 270°, and 330° on the circle when moving in clockwise direction, starting from upright position. Stimulus locations remained constant throughout the experiment, while the target appeared at each of the 6 locations with equal probability throughout the experiment, in a counterbalanced design. The search display remained visible until response.

Participants reported the tilt direction of the target, which was a T pointing either left or right with the base (long line) oriented horizontally and the head (short line) extending vertically from one end. The horizontal base was 1.5 dva in length and 0.15 dva in width, while the vertical head had equal width and was 0.75 times the length of the base, positioned at either the left or right end of the horizontal base.. Distractors shared the same color as targets but differed in shape across search conditions: In parallel search inducer trials, all distractors were straight lines (I), equal in size to the horizontal base of the target, displayed either vertically or horizontally with consistent orientation across all distractors. In serial search inducer trials, distractors were altered T shapes equal in size to target T’s, but with the vertical head off-center (25 % on stuck out one side and 65 % on other side whereas %10 was overlapping with perpendicular line). These distractors could be mirrored and rotated to 0 ° , 90 °, 180° and 270 °. In test trials, distractors were L shapes equal in size to the target, with 90 % of the short bar sticking out on one side. All distractors shared the same orientation, either vertical or horizontal, with some being mirror-reversed L shapes. These test trials were designed following (van Moorselaar & Theeuwes, 2024) to facilitate both serial and parallel search as participants could group distractors by their uniform orientation (enabling parallel search), while the high similarity between L shapes and targets could induce serial search.

Participants responded by pressing either the C or M key on a regular keyboard using the index fingers of the left and right hand to indicate the target’s tilt direction. Incorrect responses were followed by auditory feedback: a free-field auditory stimulus (330 Hz) presented via speakers for 200 ms. Gaze position was tracked and recorded throughout the experiment by a desktop-mounted Eyelink 1000 Plus with a sampling rate of 1000 Hz (one participant was inadvertently recorded at 500 Hz).

### Procedure

All participants first received verbal and written information about the experiment and then provided a written consent for voluntary participation. After cap placement and signal optimization from electrodes, at least one practice round consisting of 4 blocks was completed to teach participants how to complete the task as well as how to time blinks and eye movements. The practice section was identical to main experiment also with regards to blocked search-types but contained only 40 trials.

All participants completed 3000 trials, distributed over 5 sessions (∼35 minutes duration) that were all completed in succession on the same day. Each session contained 40 blocks with 15 trials. In each block, the first 4 trials were always inducers whereas 3 test trials were embedded in random order amongst the remaining inducers. The search type was the same throughout a block and always switched in subsequent blocks whereas the order was counterbalanced across participants. The trial flow was automated within a block whereas participants could self-determine the duration of breaks between blocks and consecutive sessions to combat fatigue.

Participants were seated in a dark room with the monitor as the only light source. After 5-dot eye-tracker calibration, participants initiated the first trial of the block by pressing SPACE bar, which was followed by the presentation of a ‘Get Ready’ message, displayed in black Arial Font for 400 ms. A blank screen with the fixation marker was presented next for 500 ms. Next, the search display was presented until response. Participants were instructed and trained to maintain their gaze fixated to the dot at the center at all times and blink when fixation dot disappeared from the screen 100 ms after response for a period ranging from 200 to 600 ms due to jitter between trials.

### EEG Acquisition and Preprocessing

The EEG was recorded from 64 pin-type, sintered, Ag-AgCl electrodes placed according to international 10-20 system. The signal was recorded at 1024 Hz via Biosemi Active 2 systems. Two external electrodes were placed on mastoids to serve as a reference offline. Additional 4 external electrodes were used to record bipolar EOG, two of which were placed below and above the right eye (Vertical) and other two lateral to the external canthi (Horizontal).

The preprocessing, filtering and epoching was handled via freely available MATLAB toolboxes, EEGLab (Delorme & Makeig, 2004) and Fieldtrip (Oostenveld et al., 2011). Offline, the data was first re-referenced to the average of mastoids and then downsampled to 500 Hz. Next, a bandpass filter (0.1 Hz highpass – 40 Hz lowpass, using pop_eegfiltnew function of EEGlab) was applied. On a bandpass filtered (1 Hz highpass – 40 Hz lowpass) copy of re-referenced continuous data, ICA was applied (using runica function with Infomax algorithm) and channel weights were extracted. We selected blink and eye-movement related components amongst 68 ICA components, via visualization of ICA maps and following the visual inspection of data before and after component removal. In the following step, trial and channel wise statistics over pre-defined window (−400 ms to 900 ms relative to search onset) were summarized to mark noisy electrodes and trials (Mean epoch variance > 1500). Upon visual inspection of all trials, noisy electrodes were replaced by spherical spline interpolation (1.20 electrodes on average) and artefacted trials were marked for exclusion from analyses (Mean = 176 trials, 5.9 % per participant). The data was epoched relative to search display onset (−400 ms to 900 ms) as well as relative to response (−700 ms – 200 ms).

## EEG Analyses

### Time-course analyses

The main analyses were conducted on 17 posterior electrodes located above visual and parietal regions (PO7, PO3, POz, Pz, P1, P3, P5, P7, Oz, P2, P4, P6, P8, O1, PO8, PO4, O2). The selection of electrodes were based on previous studies using the same analysis method (Wolff et al., 2017; Kandemir et al., 2024). The multivariate analyses were performed on independent datasets, either via training and testing on distinct conditions or by applying 8-fold cross-validation. First, the data was baseline-corrected at each electrode by removing the mean activity within 200 ms period preceding the onset of search display. Next, the data was downsampled to 250 Hz to reduce computational load.

In case of 8-fold cross-validation, we formed 8 folds of data with stratified sampling. The data in 7 folds was distributed to condition bins where the number of trials in each bin was equalized by randomly sampling the same number of trials as in the smallest bin. The data in each bin was averaged, forming a single pattern for that condition. For generalization analyses, all trials in one condition were distributed across bins and the trial-count in each bin was again equalized. A correlation-covariance matrix was estimated from the entire training set, using a custom shrinkage estimator (Ledoit & Wolf, 2004). Each trial in test set was then contrasted to the average condition patterns at each time point, quantifying dissimilarity in Mahalanobis distances (De Maesschalck et al., 2000). The output was reverse-signed to reflect similarity.

When decoding search-type, the distance difference between correct search type and the incorrect type was calculated. For conditions that contain a circular relationship (i.e. target location or target tilt), measured distances were scaled using cosine similarity and the cosine convoluted Mahalanobis distances were averaged, yielding a decoding accuracy time series in arbitrary units. This procedure was repeated 100 times to control for sampling variability and the average across repetitions was reported as the result. To improve signal-to-noise ratio the time-course data was smoothened using gaussian filter (kernel = 2 time points, 8 ms) prior to statistical analysis. The group level decoding output was tested for significance.

Temporal dynamics were investigated by assessing the generalization of neural pattern at one time point on other time points. This was either done with 8-fold cross validation when all data was from the same condition (e.g., search type or trial type) or by training on one set of trials in one condition and testing on another when assessing generalization across independent conditions. The output matrices contained decoding accuracy as the conjunction of train and test time and were assessed for statistical significance with cluster corrected sign permutation test.

EEG version of searchlight analyses (van Ede et al., 2019) was used to test the topographical generalization of signal across trial and search types. Data from each condition was prepared for decoding separately. First, signal within 300 ms time-window was averaged and this mean activity was subtracted from the data at each channel as a form of baseline. Next, data at each channel was downsampled to 100 Hz by calculating the moving average within each 10 ms window. In each round, data from each electrode and its two closest neighbours was concatenated over channel and time dimension, forming single spatio-temporal measure in each trial. Data from training set in this format was binned in accordance to distinct conditions. The trial count in each condition was equalized by randomly sampling as many trials as the trial count in the smallest bin and the data was averaged to form condition patterns. Test data was prepared the same way using the same electrodes. Each trial in test set is contrasted with averaged bins. The output was the dissimilarity of test trials relative to training conditions, calculated in Mahalanobis distance. The distance measures were reverse signed. For conditions with circular relationship, cosine-convolution was used to scale distance measures. This procedure yielded single decoding accuracy measure for each trial at each electrode, reflecting generalization of neural pattern across trial or conditions. The mean decoding accuracy over trials and subjects was contrasted against zero with a sign permutation test and statistically significant results indicated electrodes where signal significantly generalized across conditions. Topographical plots are used to reflect decoding accuracy / generalization with bright colors indicating stronger effects.

In order to control for the role of eye movements on decoding, we decoded search type and item location from gaze position recorded from an eyelink 1000 eye-tracker. Trial-count for EEG and eye tracker showed small difference because the eye-tracker sometimes missed eye position. Therefore, we also decoded EEG using the same trials and assessed the correlation of these measures. Correlation assessment had three versions to ensure EEG decoding did not stem from gaze position or movement. First we assessed correlation at each time point across trials and tested Fisher transformed correlation coefficients against zero with cluster corrected permutation test. Absence of time-course correlation could be caused by a lag in the emergence of the effect in EEG after stemming from the eye. Therefore, we averaged decoding accuracy from eye and EEG within 100 – 700 ms period and correlated these two then tested the Fisher transformed correlation coefficients against zero using group permutation test. Finally, we calculated mean decoding from eye and EEG recordings at subject level and assessed correlation against zero to see if decoding was high in participants with stronger eye decoding.

### Correlation analysis

In order to assess link between search types and response time in test trials, we assessed the generalization of target location between inducers with distinct search types and test trials. For each trial we had two decoding results, each showing generalization to one type of search inducer. We temporally averaged time-course signal within 100-300 ms period since this period was the only window where generalization differed in test trials with both search type. We excluded trials that were slower than 3 standard deviation of mean RT and correlated trialwise decoding accuracy difference and RT. Extracted correlation coefficients for each subject were Fisher transformed and group mean was tested against zero with t-test.

### Statistical assessment

The time-course decoding and temporal generalization cluster corrected non-parametric permutation test was used. A null distribution was first formed by alternating the sign of the data at each time point randomly with 0.5 probability (N _perm_= 100 000). The average decoding was then compared to null distribution and the proportion corresponding to the mean decoding accuracy averaged over trials and subjects was reported as p value . For time-course data, a cluster correction with 0.05 as cut-off was applied to control for false-positives (Spaak, 2015). Additionally for temporal generalization analyses, where data trained at one time point was contrasted with data across all time points, we employed another test to assess dynamicity of the signal. Here, signal along the diagonal would indicate time-specific decoding whereas vertical and horizontal extensions would reflect generalization into preceding and/or subsequent time points. Here, we assessed whether diagonal was significantly larger than the average of horizontal and vertical time points by using a custom function (Myers et al., 2015). Areas that were significantly lower than diagonal were marked.

When decoding was conducted over a time-window using 17 posterior electrodes or a subset of electrodes (e.g. Searchlight analysis), yielding a decoding accuracy measure for each trial, mean decoding over subjects was tested with non-parametric group permutation test against zero. For this, the sign of data was randomly alternated with 0.5 probability for 10000 times to form a null distribution. The proportion corresponding to actual data was reported as the p value. All tests were two-tailed unless explicitly stated.

### Behavioural analysis

Response time and accuracy were primary behavioural measures. Mean accuracy under different trial conditions (Inducer and Test) as well as search conditions (Serial and Parallel) were calculated separately for each participant and Repeated Measures ANOVA (RM-ANOVA) was tested for two main effects and their interaction using JASP (JASP Team, 2024). For RT analysis, we first filtered out trials that were slower than 200 ms, considering these responses as reflexive rather than planned responses. Next, we excluded outliers by removing trials with responses slower or faster than 3 standard deviations above/below the mean reaction time for each subject. Reaction times in remaining trials were averaged for different trial and search types. The main effects of search type and trial condition and their interaction was assessed with RM-ANOVA using JASP.

## Data Availability

Processed and anonymized EEG and behavioural data reported in this manuscript, as well as MATLAB scripts and their output can be accessed via this link: (https://osf.io/qg9b2/overview?view_only=eb6019c1026d4dafab52efb1212fb6a1)

## Acknowledgements

This project was funded by the European Research Council (ERC; grant 833029-LEARNATTEND), and the Dutch Research Council (NWO; grant 406.21.GO.034) in grants awarded to J. Theeuwes. We thank Anna Sophia Rauleder for her help during data collection.

---

Supplements – I

EEG and MEG decoding is vulnerable to confounding by gaze position and micro-saccades (Mostert et al., 2018). We decoded search type using gaze position recorded by eye-tracker. Decoding search type yielded significant effects briefly after the onset of search display and which stronger after 400 ms (Fig. S1A – top panel, Search type (blue), 64 ms – 900 ms relative to search onset, *p* < 0.001). Decoding EEG from 17 posterior electrodes using the same trials also yielded significant decoding (Fig. S1A – top panel, Search type (black), 32 ms – 900 ms relative to search onset, *p* < 0.001). To assess if these measures were linked, decoding accuracy from Eye and EEG data was correlated at each time point per subject and time-course Fisher transformed correlation coefficients were tested for significance. Cluster-corrected sign-permutation test yielded no significant clusters with correlation above zero (Fig. S1A – bottom panel, black). The link between gaze position and EEG could take place with a time-lag, preventing us from detecting their relationship. Therefore, we averaged decoding outputs over time dimension within 100- 700 ms period relative to search onset. Fisher transformed correlation coefficients did not significantly differ from zero (Fig. S1A –left, boxplot for Fisher transformed correlation coefficients, blue rectangle marks time-zone of correlation). Finally we tested correlation at subject level, by correlating mean decoding within 100-700 ms window using Eye and EEG data (Fig. S1B). This tested whether EEG decoding was stronger in participants with strong decoding from gaze position. Although a positive trend was observed, this was not statistically significant (*R^2^* = 0.06; *p* = 0.24). We conclude that of search type decoding in EEG is not driven by the variation in gaze position.

**Figure S1.**
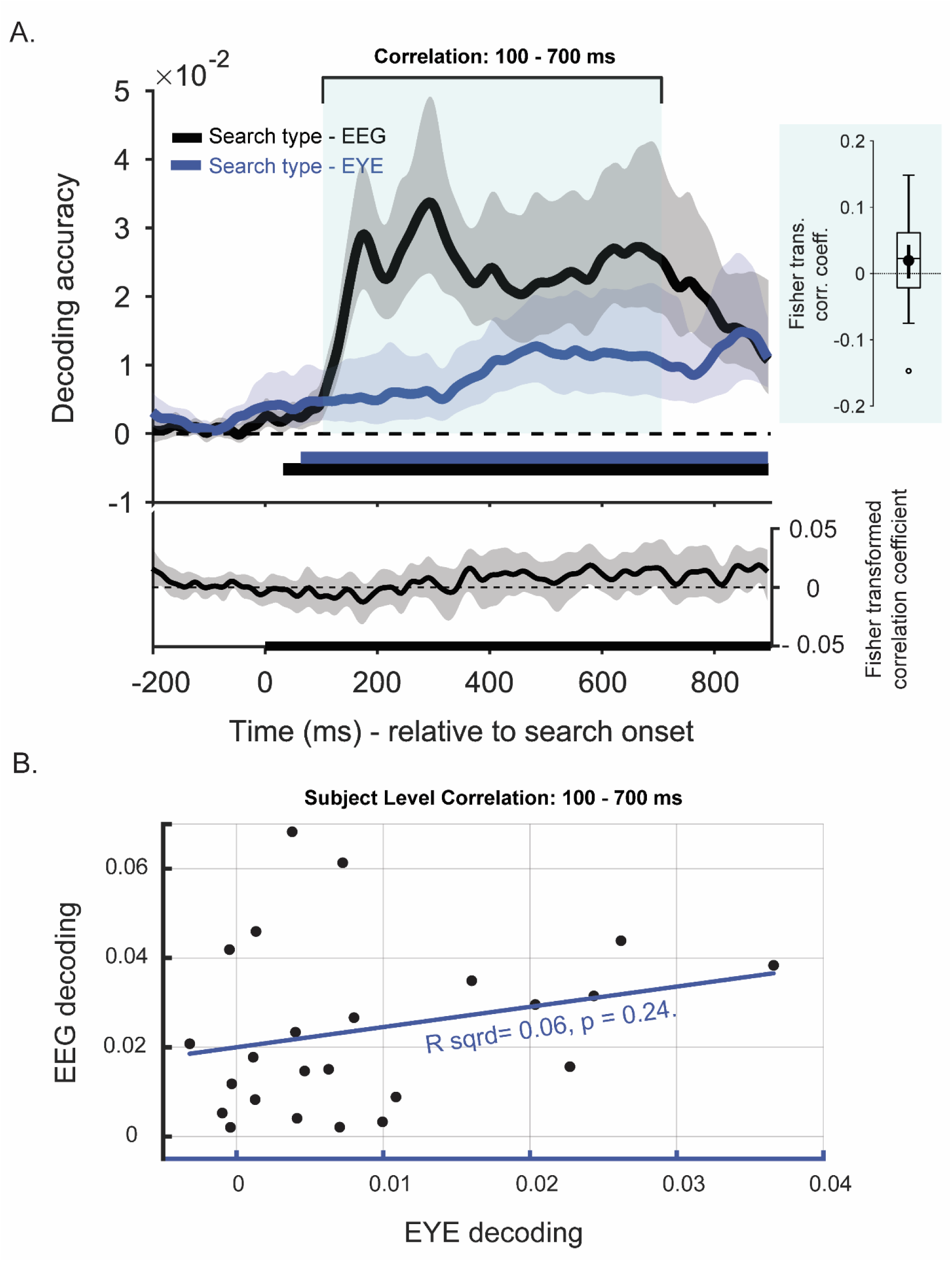
Decoding accuracy for search type using EEG and EYE data, the correlation of decoding accuracy at each time point as a function of time and across trials within 100-700 ms window (A), and across subjects within the same 100-700 ms window (B). *Note.* Assessing the contribution of gaze position to decoding. **A-Top panel.** Time-course decoding of search type using EEG (black) and eye data (blue). Solid lines reflect mean decoding accuracy as a function of time, indicating generalization of location modulation across inducer and test trials in specific conditions. Shades around solid lines mark 95 % C.I. Solid bars below baseline mark the time-window with statistically significant decoding according to cluster-based permutation test (*p* < 0.05, two-tailed). **A-Bottom panel.** Time-course Fisher transformed correlation coefficients reflecting relationship between Eye and EEG decoding. **A-Left panel**. Boxplot for Fisher transformed correlation coefficients for decoding from Eye and EEG decoding averaged over 100 – 700 ms window, also marked in time-course with blue shaded zone. Dot in the middle marks the mean where solid line indicates median. Small horizontal bar shows 95 % C/I and the whiskers of the box reflect 1.5 IQR. B. Scatterplot showing subject wise mean decoding from Eye (x-axis) and EEG (y-axis) with a line fitted and correlation coefficient tested for significance.

Supplements – 2

The location of target was also traceable from gaze position both when test trials were in serial search (Fig. S2A – top panel, Target loc- Serial EYE, blue, 348 ms – 652 ms relative to search onset, *p* = 0.003) and parallel search condition (Fig. S2A – top panel, Target loc-Parallel EYE, purple, 304 ms – 892 ms relative to search onset, *p* < 0.001). The same trials yielded significant decoding using EEG when search was serial (Fig. S2A – top panel, Target loc- Serial EEG, cyan, 248 ms – 780 ms relative to search onset, *p* < 0.001) and parallel (Fig. S2A – top panel, Target loc- Parallel EEG, fuchsia, 136 ms – 824 ms relative to search onset, p < 0.001). Although the differences in onset times suggested that processing of target location was more likely inducing gaze shifts, we assessed correlation between these two measures. Cluster-corrected sign-permutation test yielded no significant clusters for correlation between target location decoded from Eye data and EEG neither during parallel nor serial search (Fig. S2A – bottom panel, cyan and fuchsia). Correlation across trials when averaging over 100-700 ms window was also not significant. We also failed to find any correlation at subject level (*R^2^*-Parallel = 0.03; *p* = 0.46; *R^2^*- Serial = 0.06; *p* = 0.24). We conclude that of search type decoding in EEG is not driven by the variation in gaze position.

**Figure S2.**
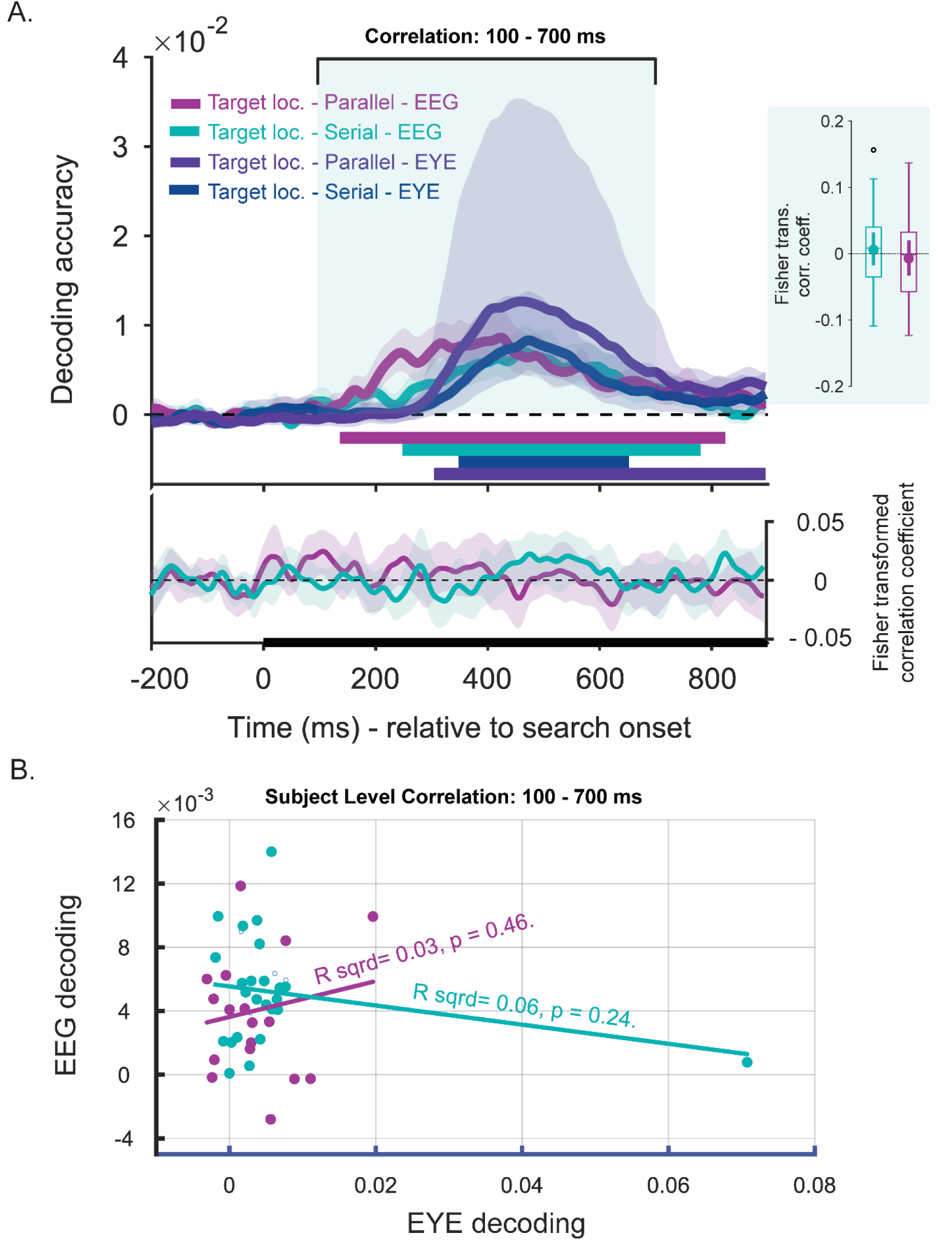
Decoding accuracy for target location in distinct search conditions using EEG and EYE data collected during test trials, the correlation of decoding accuracy at each time point as a function of time and across trials within 100-700 ms window (A), and across subjects within the same 100-700 ms window (B). *Note.* Assessing the contribution of gaze position to decoding of target location in test trials with serial and parallel search. **A-Top panel.** Time-course decoding of search type using EEG in parallel (fuchsia) and serial (cyan) search as well as using eye data in serial (blue) and parallel search (purple). Solid lines reflect mean decoding accuracy as a function of time, indicating generalization of location modulation across inducer and test trials in specific conditions. Shades around solid lines mark 95 % C.I. Solid bars below baseline mark the time-window with statistically significant decoding according to cluster-based permutation test (*p* < 0.05, two-tailed). **A-Bottom panel.** Time-course Fisher transformed correlation coefficients reflecting relationship between Eye and EEG decoding during test trials with serial (cyan) and parallel search (fuchsia). **A-Left panel**. Boxplot for Fisher transformed correlation coefficients for decoding target location in different search conditions using Eye and EEG data, averaged over 100 – 700 ms window, also marked in time-course with green shaded zone. Dot in the middle marks the mean where solid line indicates median. Small horizontal bar shows 95 % C/I and the whiskers of the box reflect 1.5 IQR. B. Scatterplot showing subject wise mean decoding from Eye (x-axis) and EEG (y-axis) with a line fitted and correlation coefficient tested for significance.

Supplements – 3

**Figure S3.**
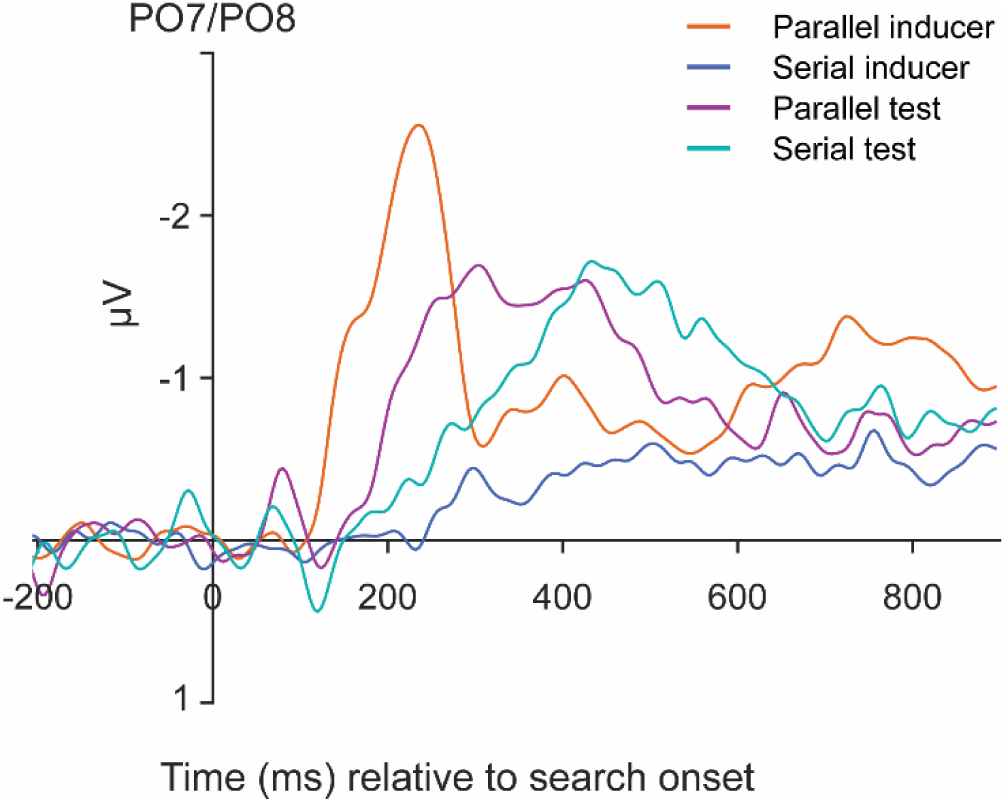
Decoding accuracy for target tilt direction separately for distinct locations in parallel (A) and in serial inducer trials (B). *Note.* Event-related potentials showing contra-ipsi difference over PO7 and PO8, averaged separately over Parallel inducer, Serial inducer, Parallel test and Serial test trials, aligned to search display onset. Data baselined to −200 ms relative to search onset, presented with Y axis reversed.

Figure S3 presents contra-ipsi difference waves calculated over PO7 and PO8 electrodes, separately in each search condition across trial types. The N2Pc component, associated with the shifting of attention towards the target (Eimer, 1996) is usually present between 100-300 ms window. Here, we clearly see a large peak during parallel inducer trials, approximately by 240 ms. A delayed N2Pc emerges for Parallel test trials at 300 ms and for Serial test trials at 430 ms. For serial inducers, earliest elevation is observed around 300 ms although signal peaks at around 750 ms. These findings indicate that the timing and magnitude of attentional selection differed across search modes, with earlier and stronger N2pc responses during parallel search. The time course of ERPs for test trials aligns with target location decoding presented in Figure 2B.

Supplements – 4

Parallel and serial search differ with regards to all or some of items being attended within the same time window. Our design allowed us to check if items at all locations were processed at the same time in parallel inducer trials. When we decoded our data for target item tilt, we observed that decoding accuracy did not significantly differ across distinct locations and the time window associated with significant decoding largely overlapped (Fig. S4A – Location 1, fucsia, 224 ms – 720 ms relative to search onset, *p* < 0.001; Location 2, red, 260 ms – 736 ms and 752 – 900 ms relative to search onset, *p* < 0.001 and p =0.028, respectively; Location 3, burgundy, 256 ms – 772 ms relative to search onset, *p* < 0.001; Location 4, violet, 236 ms – 900 ms relative to search onset, *p* < 0.001; Location 5, blue, 228 ms – 760 ms relative to search onset, *p* < 0.001; Location 6, purple, 260 ms – 608 ms and 624 – 736 ms relative to search onset, *p* < 0.001 and p = 0.04, respectively, all *two tailed*). Although cluster corrected tests do not reliably test for onset times (Sassaghanen & Draskow, 2019), the absence of significant difference in the strength of decoding and the overlapping of all 6 locations provides some evidence that all locations were attended at the same time.

During serial search, target tilt was traceable with a statistically significant strength only in location 2 and 5 (lateral positions) which emerged approximately 80 ms later in time relative to parallel search (Fig. S4B., Location 2, red, 334 ms – 900 ms relative to search onset, *p* < 0.001; and, Location 5, blue, 376 ms – 732 ms relative to search onset, *p* = 0.002, , both *two tailed*). Target tilt was not traceable with statistical significance at other locations indicating that items at these locations were attended at varied time points). These results clearly show differences in target processing during serial and parallel search.

**Figure S4.**
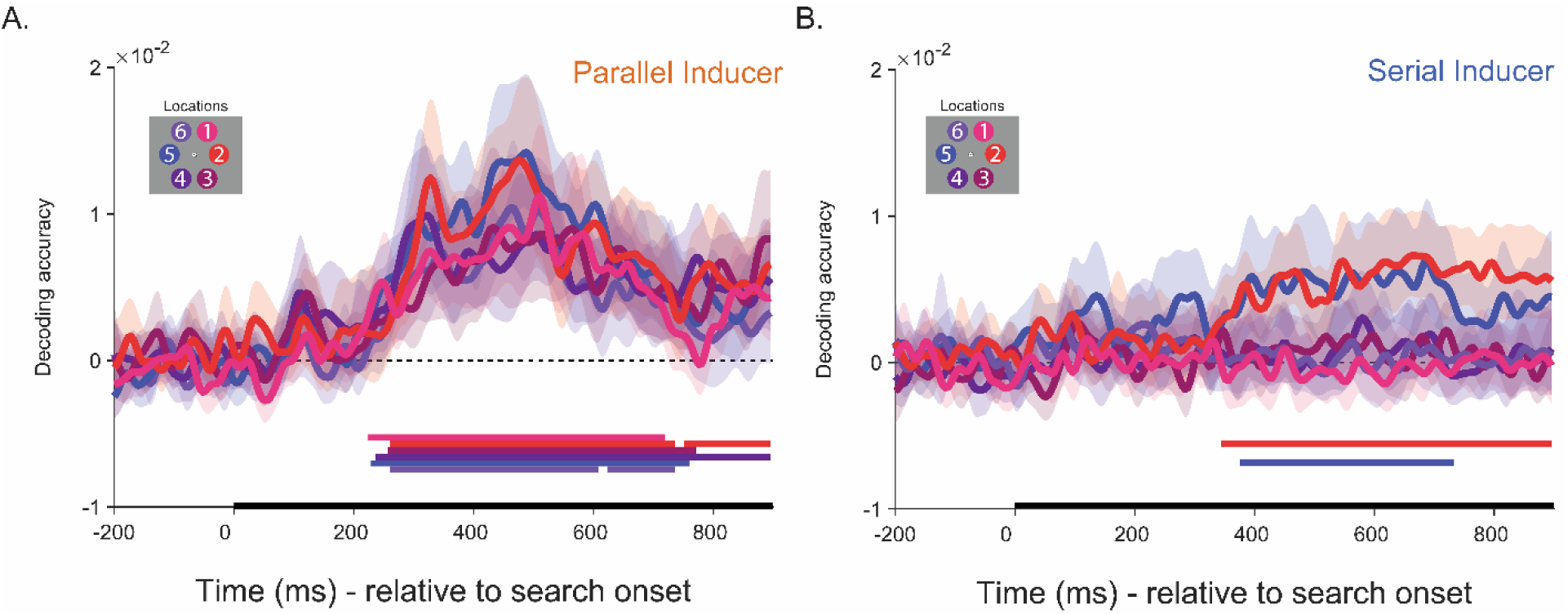
Decoding accuracy for target tilt direction separately for distinct locations in parallel (A) and in serial inducer trials (B). *Note.* Time-course decoding of target tilt direction at each location during parallel (A) and serial (B) inducer trials, reflecting that all locations are processed at the same time during parallel search but at distinct times during serial search. (A-B). Solid lines reflect mean decoding accuracy as a function of time, indicating task relevant tilt (left vs. right) of target T. Each color represents a different location. Shades around solid lines mark 95 % C.I. Solid bars below baseline mark the time-window with statistically significant decoding according to cluster-based permutation test (*p* < 0.05, two-tailed).

Supplements – 5

To contrast the deployment of attention during distinct search conditions, we assessed the generalization of the target location representation across search modes in inducer trials (Fig. S5). Specifically, a cross generalization of location decoding revealed similar patterns between two search modes, indicating a general attentional spotlight was used in both search modes. Importantly, this spatial orienting was extremely consistent in parallel search, reflecting that the initial deployment of attention often landed on the search target. However, the temporal characteristics differed markedly: In parallel inducers, spatial attention was deployed consistently and early, typically landing on the target location, whereas in serial inducers this localized attention component was temporally variable and distributed across a longer timescale, reflecting trial-by-trial variability in when the attentional spotlight located the target.

**Figure S5.**
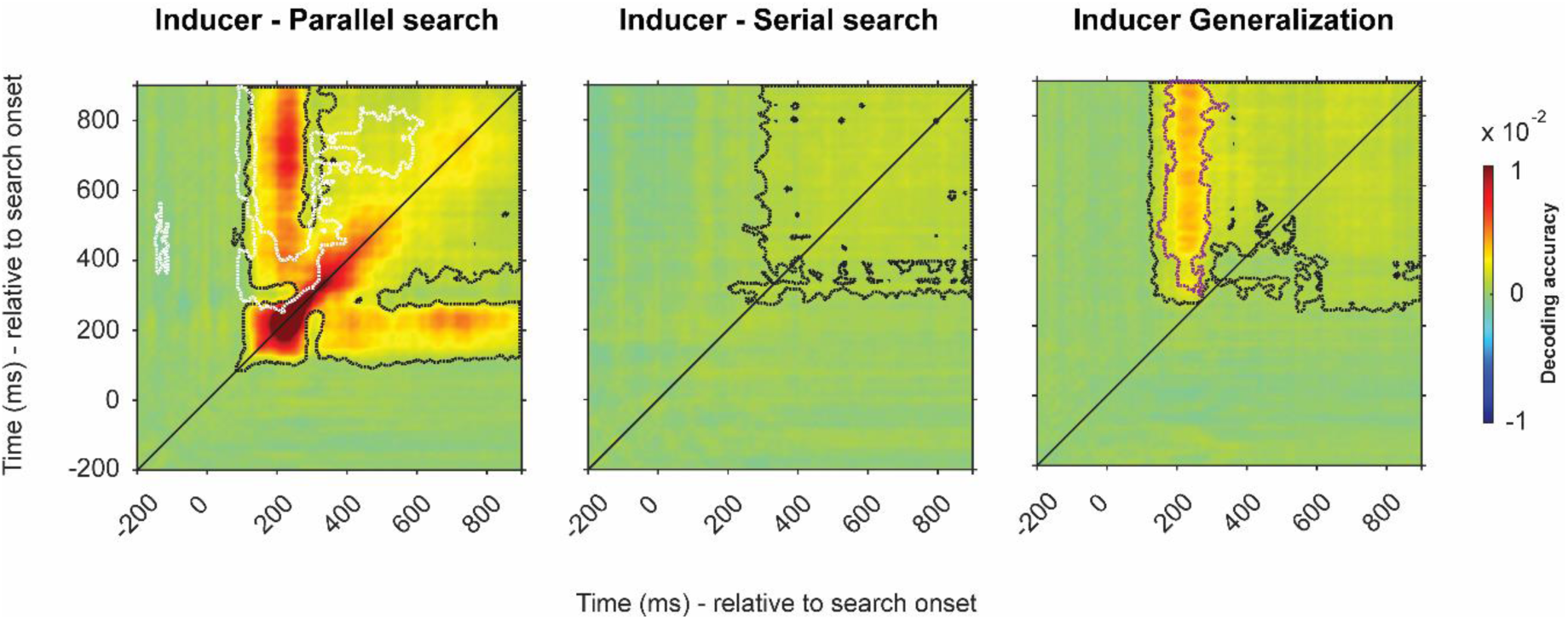
Temporal generalization of neural code for target location in parallel (left), serial search inducers (middle) and the generalization across (right), relative to search onset time. *Note.* Decoding target location by training at a time point and testing on all time points. Temporal generalization is assessed within and across each search type in inducer trials. Dotted black lines mark clusters with statistically significant decoding against zero (*p* < 0.05, two-tailed). Dotted white lines mark clusters with diagonal decoding is significantly higher than the average of vertical and horizontal, indicating dynamic decoding period whereas dotted red lines mark clusters with diagonal is significantly lower than vertical and horizontal, indicating reactivation (*p* < 0.05, one-tailed).

